## Supplementary Information for "Broadscale dampening of uncertainty adjustment in the aging brain"

**This PDF file includes:**

Supplementary Figures S1 to S6
Supplementary Text S1 to S7
Supplementary Table S1-5
Supplementary References

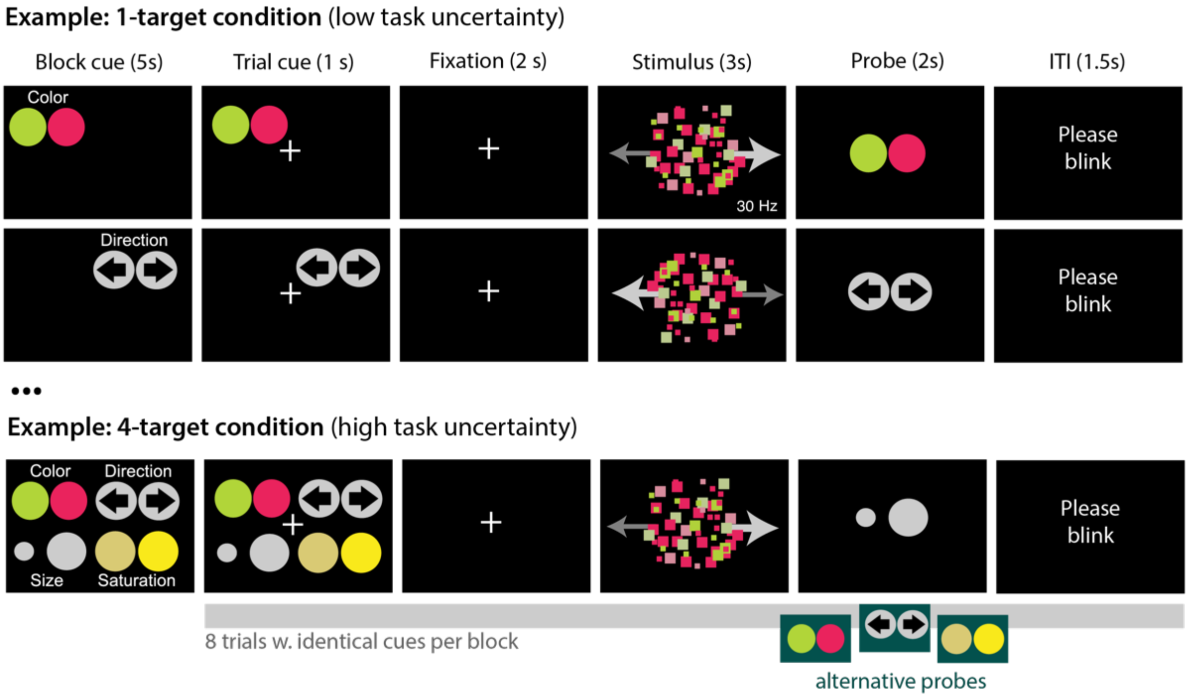


**Figure S1-0. Examples of design phases.**

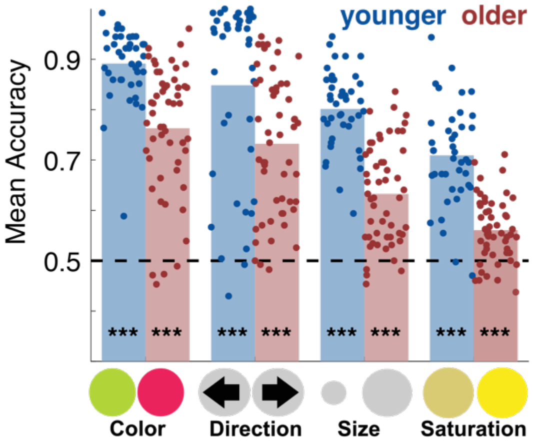


**Figure S1-1. Average accuracy across uncertainty conditions.** Younger and older adults on average performed the task above chance for all features that were probed. Statistics are based on one sample t-tests against chance level (0.5 in this 2AFC task). *** $p < .001$.

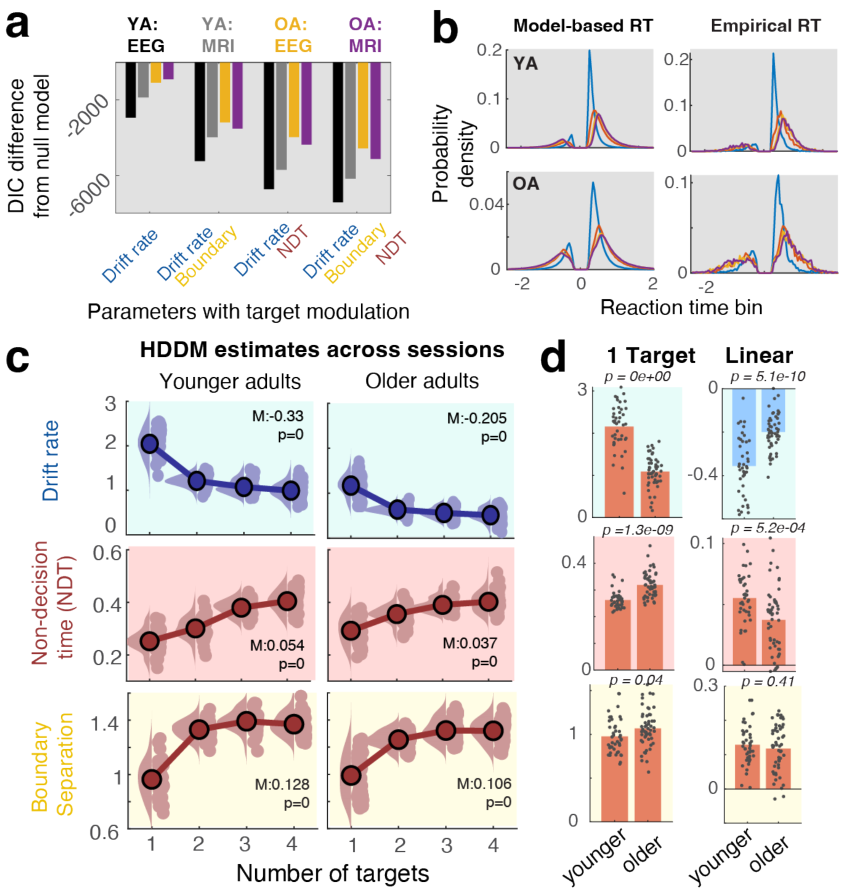


**Figure S1-2. Age-related uncertainty adjustments to decision processes.** (a) DIC-based model comparison indicates that a model, including uncertainty modulation of drift rates, non-decision times, and boundary separation provides the best group fit to the behavioral data. (b) Posterior predictive checks for the full model (shown for the EEG session). Negative RTs indicate incorrect responses. Model-based ("posterior predictive") values were sampled 50 times within each participant and condition (as implemented in the HDDM package), and probability density (100 RT bins) was estimated first within-subject across all samples, and then averaged across participants. In empirical data, probability densities were estimated across all participants due to the sparse within-subject RT counts. (c) Uncertainty modulation of HDD parameter estimates, averaged across sessions. Statistics refer to paired t-tests of linear slopes against zero. Data are within-subject centered. (d) Age comparison of single-target parameter estimates (left) and linear uncertainty effects (~age x target load interaction). Statistics refer to unpaired t-tests.

**Text 1-2. Uncertainty and age effects on non-decision time and boundary separation.** The main analyses targeted drift rate as the main parameter of interest. Given that the best-fitting model (Figure S1-2ab) included uncertainty variation also for non-decision times as well as boundary separation, we explored the potential variation of the latter two parameters with age and uncertainty (Figure S1-2c). In contrast with younger adults, older adults had significantly longer non-decision times, and larger boundary separation, suggesting that more evidence was collected prior to committing to a choice. There is some evidence from 2AFC tasks that older adults adopt decision boundaries that are wider than the boundaries of younger adults (Starns & Ratcliff, 2010, 2012) [but see (McGovern et al., 2018)], which may signify increased response caution . In both age groups, we observed uncertainty-related increases in non-decision times, albeit more constrained in older adults, as well as similar increases in boundary separation as a function of rising uncertainty (see Figure S1-2d). Notably, the uncertainty effect on boundary separation was not consistently reproduced by either the integration threshold of the domain-general CPP (Figure S1-5b), or the effector-specific contralateral beta power threshold (Figure S1-6d), highlighting uncertainty regarding the true effect on behavioral response caution, or neural proxy signatures thereof. These discrepancies deserve further attention in future work and may suggest that a model with alternative parameter constellations could provide a more coherent description. Convergence of the current model with our previous results in younger adults (Kosciessa et al., 2021) ultimately argues for robust drift rate inferences that were independent from the specific model choice**.**

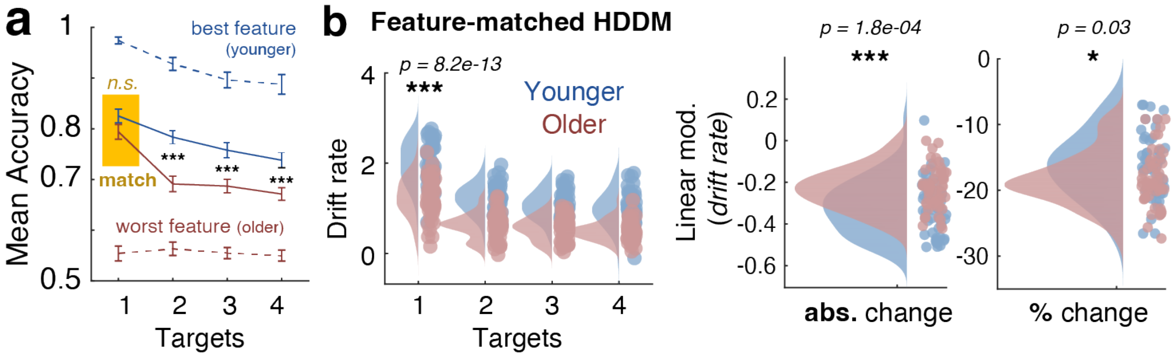


**Figure S1-3. Drift rate differences do not arise from accuracy ceiling or floor effects.** (**a**) Difficulty with selectively distinguishing individual features is a major component of the task, which may contribute to age differences in single-target drift rates. While on average, younger adults' single-target responses were more accurate than those of older adults, this differed between features (see Fig. S1-1). Excluding the most accurate ('best') feature for younger adults and the least accurate ('worst') feature for older adults matched groups regarding their accuracy in the single-target condition. In this scenario, older adults showed more pronounced accuracy decreases under uncertainty, relative to younger adults. Data are means +/- SEs and include data from EEG and fMRI sessions. n.s.: p = 0.13; ***1: p = 2.6e-05; ***2: p = 6.1e-04; ***3: p = 8.2e-04. (**b**) Drift rate estimates for an HDDM model that only included age-matched features ("match" in ***a***). This model indicated retained age differences in single-target drift rate and absolute drift changes under uncertainty (right). Older (vs. younger) adults showed stronger *relative* drift rate reductions from the single-target baseline.

**Text S1-3a. Drift rate effects for accuracy-matched features.** Our analysis indicated that older adults on average showed reduced behavioral uncertainty costs. However, these uncertainty costs are thought to arise from attending to a varied feature set, whose discrimination also varies between age groups when only a single feature is relevant. To examine whether potential ceiling or floor effects in feature-specific accuracy (e.g., due to varying perceptual uncertainty) acts as a between-group confound, we sorted features according to their single-target accuracy in each participant, and averaged accuracy according to such "preference" within each age group. This revealed that three out of the four features elicited comparable single-target accuracy between age groups, whereas only the best feature of younger adults, and the worst feature of older adults could not be matched (Figure S1-3a). To test the robustness of unmatched drift rate estimates (Fig. 1b), we created HDDM models that excluded the most preferred feature of younger adults, and the individually least preferred feature in older adults (i.e., only including "matched" features). Results from this control analysis are shown in Figure S1-3b. We observed retained age differences in single-target drift rates, as well as uncertainty-related drift rate changes that mirrored our main results. These results indicate that baseline feature differences are likely not the principal origin of age and uncertainty drift rate differences.

**Text S1-3b. More pronounced relative performance decreases in older adults.** Compared with younger adults, older adults' drift rates were lower across levels of target load (Fig. 1b). To test whether drift rates across all set sizes show similar proportional age changes, we calculated relative drift rate changes. Arguing against uncertainty-independent age differences in drift rate, we observed larger *relative* drift rate decreases under uncertainty in older as compared with younger adults (see Fig. S1-3b right for feature-matched HDDM; similar results were obtained in the main model). This indicates that despite being smaller in absolute terms, older as compared to younger adults suffered stronger *relative* drift rate losses once uncertainty was introduced. This mirrored larger accuracy decreases in matched features once uncertainty was introduced (Fig. S1-3a). Taken together, this indicates that uncertain multi-tasks contexts present an outsized challenge to older adults' performance, over and above challenges in single-target specificity (Jaroslawska et al., 2021). For our main analyses that target inter-individual relations, we focus on absolute uncertainty-related drift rate changes due to their relation to neural uncertainty adjustment in prior work (Kosciessa et al., 2021), and the computational interpretability of absolute drift rates at each target load.

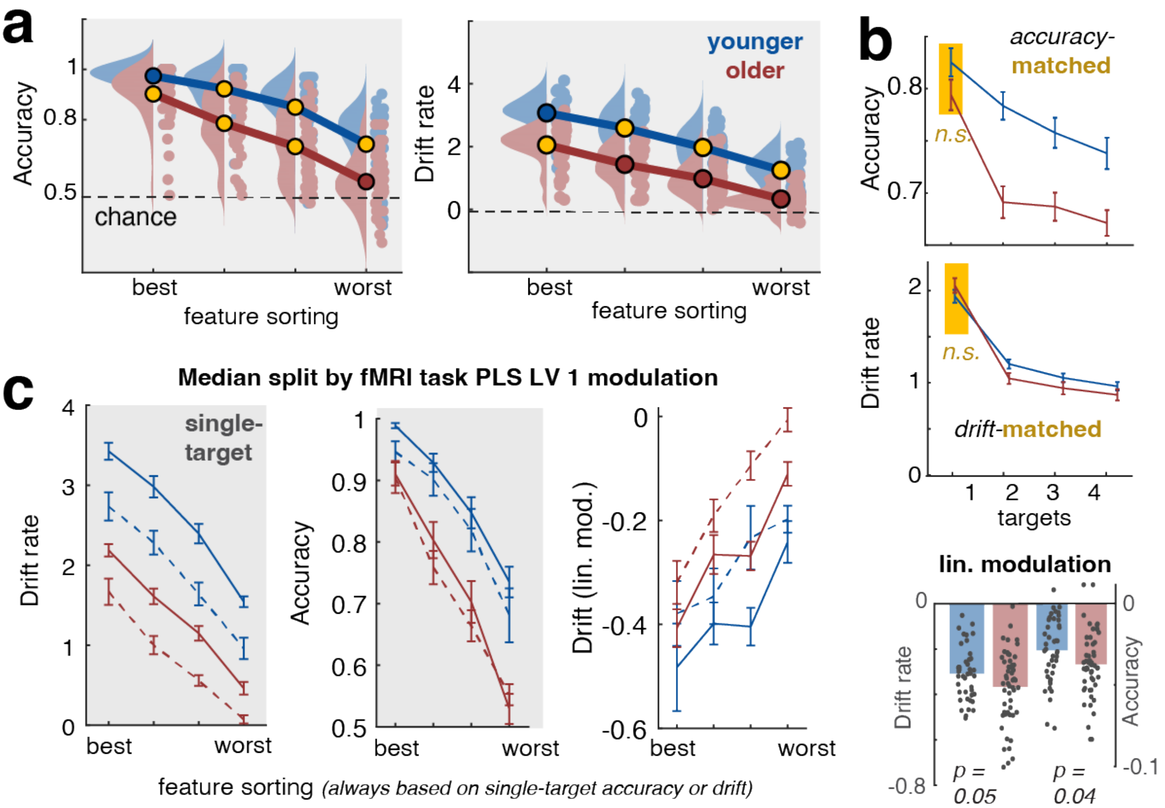


**Fig. S1-4. Exploration of feature-specific accuracy and drift rate variation.** (a) Single-target accuracy and drift rate for features that are sorted by decreasing single-target accuracy and drift rate. Yellow dots indicate the conditions selected for matching in b. (b) Matching based on single-target accuracies and drift rates indicates stronger *relative* performance decreases with uncertainty in older compared with adults. (c) A median split of fMRI Brainscores (trichotomized split | full lines: largest BOLD mod.; dashed lines: smallest BOLD mod.; cf. Fig. S6-3) indicates that larger neural modulation is inter-individually and group-wise linked to higher single-trial drift rates (but not differential accuracy; see also Fig. S6-3), independent of feature preference. This suggests that neural modulation is not primarily constrained due to varying choice difficulty for individual features, but rather by a global operation (e.g., cue-guided selectivity, distractor suppression, etc.) that is shared across features. Means +- SEMs.

**Text S1-4. Exploration of feature-specific accuracy and drift rate variation.** We performed exploratory analyses to further characterize behavior (accuracy, rt) and drift rate as a function of probed feature. To this end, we estimated all canonical HDDM parameters also for variations in the feature that has been probed. We caution that the large parameter space may lead to some convergence issues, which is why the main model does not estimate parameters for separate features. We sorted single-target accuracy and drift rate in descending order (Fig. S1-4a). Like accuracy, selecting features such that single-target drift rates are matching indicates *larger* uncertainty-induced drift rate decreases (Fig. R1-4b, see also Text S1-3b). However, it is debatable whether this is a *reasonable* comparison particularly for drift rates: an analysis that matches "selective" performance (i.e., drift rates under low uncertainty) between age groups also removes key age differences in neural engagement. Namely, we observed a closer link between within-group fMRI modulation and single-target drift rates, but not accuracy (Fig. R1-4c**,** see also S6-3). This was independent of which feature was probed, and thus unrelated to varying perceptual uncertainty between features. This reinforces the notion that perceptual uncertainty per se (which in psychophysics is sometimes titrated to comparable *accuracy* levels) is not the principal predictor of the constrained neural uncertainty modulation either within or between age groups.

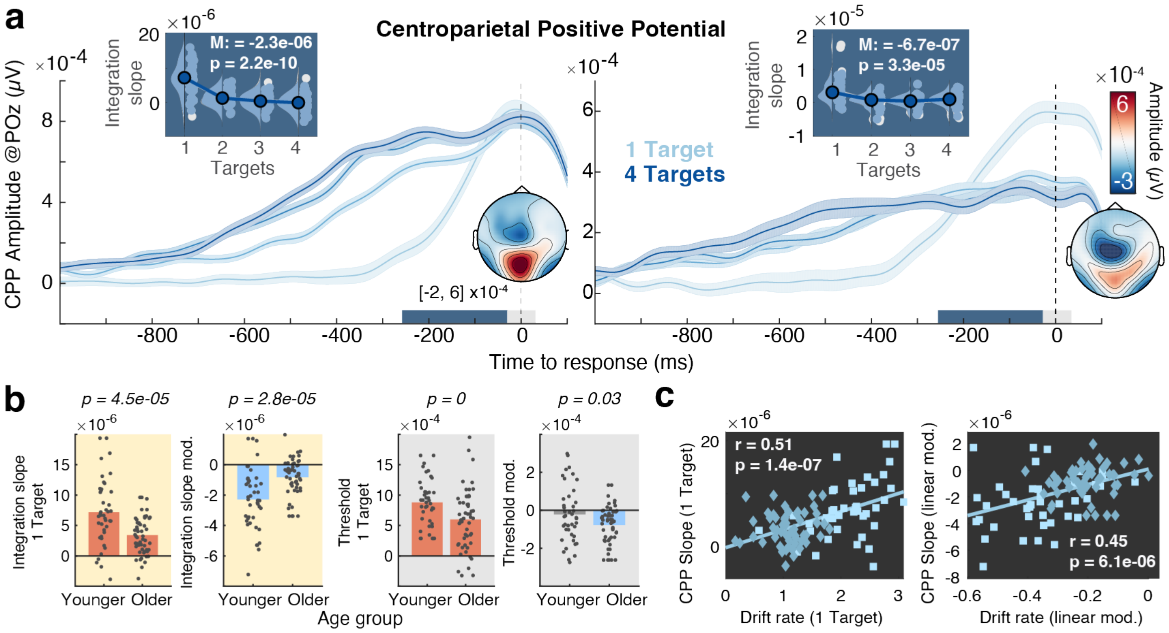


**Figure S1-5. Centroparietal Positive Potential (CPP) as a signature of domain-general evidence integration.** (a) Modulation of CPP as a neural signature of evidence accumulation (mean ± within-subject SEM). The integration slope of the response-locked CPP decreases with increasing uncertainty. Traces are mean ± within-subject SEM. Insets show CPP slope estimates from −250 to −50 ms relative to response execution. (b) Age comparison of CPP integration slopes (yellow background) and CPP integration thresholds (grey backgrounds). (c) CPP estimates of evidence integration converge with behavioral drift rate estimates at the interindividual level, both w.r.t the single-target condition (r(93) = 0.45, 95%CI = [0.27,0.59], p = 6.1e-06; *age-partialed*: r(92) = 0.27, 95%CI = [0.08,0.45] p = 0.01) and linear effects of target number (r(93) = 0.51, 95%CI = [0.34,0.64], p = 1.4e-07; *age-partialed*: r(92) = 0.34, 95%CI = [0.14,0.5] p = 9.3e-04).

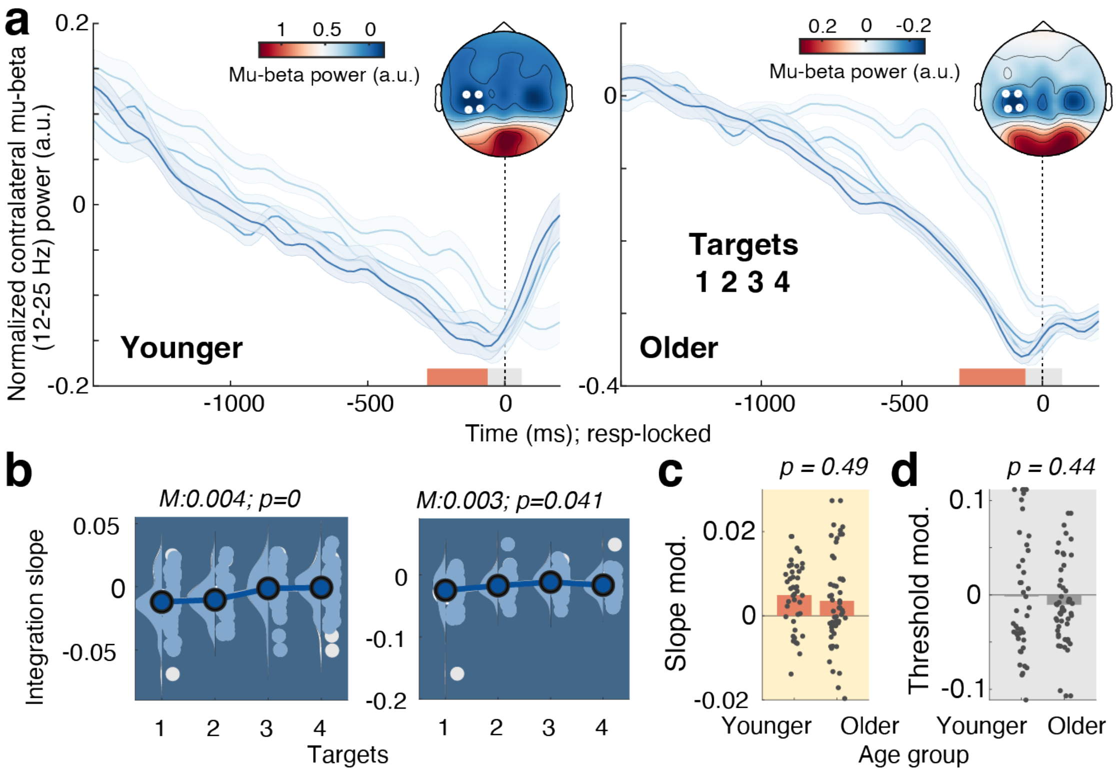


**Figure S1-6. Contralateral beta power as a signature of motor-specific response preparation.** (a) Pre-response desynchronization of contralateral mu-beta power shallow with increasing number of targets. Traces show means +/- within-subject SEM. (b) Linear slope estimates, estimated via linear regression from -250 ms to -50 ms, relative to response. Data are within-subject centered for visualization. Statistics refer to paired t-tests of linear slopes against zero. (c, d) Age comparison of linear modulation of beta slopes (c) and integration thresholds (d) by target load. Statistics refer to unpaired t-tests.

**Text S1-6. Motor-specific response preparation.** In addition to the domain-general CPP, we also investigated motor-specific contralateral beta power (Figure S1-6a). Extending results from behavioral modeling, and CPP integration slopes, we observed a shallowing of pre-response beta power build-up, suggesting decreases in response preparation (Figure S1-6b). However, such shallowing was not statistically different between age groups (Figure S1-6b), thus deviating from the age x load interaction that we observed for the remaining integration signatures. Furthermore, linear changes in beta slope as a function of target load were neither associated with linear drift changes (r(93) = -0.03, 95%CI = [-0.23,0.17], p = 0.77) nor CPP slopes (r(93) = -0.11, 95%CI = [-0.3,0.09], p = 0.29) across age groups. The parameters were also not directly related in the single-target condition (*drift rates*: r(93) = 0.18, 95%CI = [-0.02,0.37], p = 0.07; *CPP slopes*: r(93) = -0.06, 95%CI = [-0.26,0.14], p = 0.55). Motor-specific response preparation thus appears to partially dissociate from effector-unspecific evidence integration at the individual level.

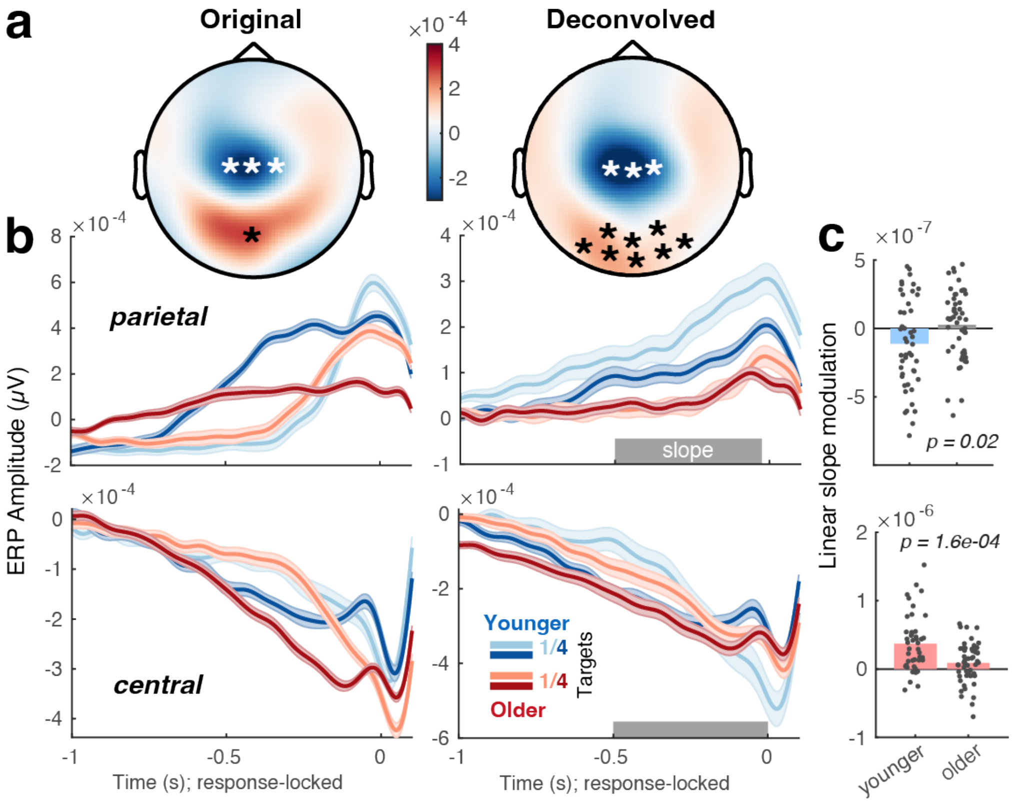


**Supplementary Fig. S1-7. Deconvolution analyses for response-locked potentials.** (a) Average peri-response topographies from -250 ms to 100 ms before (left, cf. Fig. 1c) and after (right) probe onset deconvolution. Black stars indicate sensors selected for parietal/CPP potentials, whereas white stars indicate central channels selected for motor-related potentials. (b) Response-related potentials before (left) and after (right) probe onset deconvolution. Means +- within-subject SEM. Parietal (top) and central (bottom) potentials correspond to the channels indicated in a. (c) Uncertainty modulation (linear) of pre-response slopes (deconvolved data; cf. Fig. 1c) by age group. Pre-response slopes were estimated during the time windows indicated by the shaded areas in b.

**Supplementary Text S1-7. Deconvolution of probe-related from response-locked ERP potentials.** Recent work proposes that pre-response CPP ramping could be sufficiently explained by a probe-related appraisal process – rather than choice – that is jittered in time due to response speed variations (Frömer et al., 2024). To explore this here, we conducted deconvolution analyses using the unfold toolbox (Ehinger & Dimigen, 2019) implemented in MATLAB. By providing event-specific regression formulas, this approach can disentangle temporally overlapping ERP responses. We jointly modelled probe onset and response regressors using FIR basis functions, time expanded +/- 1 seconds around the respective events (Frömer et al., 2024). Data were 8 Hz lowpass-filtered, no baseline correction was applied, amplitudes > 250 μV were removed. Consistent with the CPP reflecting a stimulus-related P300 potential (O'Connell et al., 2012; Twomey et al., 2015), posterior topographies changed following removal of onset-related activity (Fig. *S1-7*). However, parieto-occipital potentials more exhibited residual ramping that peaked slightly prior to response onset, in line with a choice-related process. Age differences in the uncertainty modulation of residual slopes (linear fits: -500 to -50 ms) resembled those in the main analysis (cf. Fig. 1c), albeit with no uncertainty-related slope adjustment observed in older adults. Analogous to beta power dynamics (Fig. *S1-6*), a central motor-related potential indicated that uncertainty shallowed slopes of the deconvolved motor negativity (linear fits: -500 to -0 ms).

While onset deconvolution captures the impact of probe processing on response-locked potentials, it provides inconclusive information regarding an accumulation-to-bound decision process. Differences in evidence integration rate can emerge rapidly after probe onset, persist until response (see e.g., Churchland et al., 2008; Twomey et al., 2015), and produce differential RT distributions (Twomey et al., 2015). In the face of temporal covariance, considering unique response-locked variance would remove substantial "choice" signal of interest. In the face of this ambiguity regarding the latent processes contributing to observed potentials, multiple observations support a choice-related account here: First, CPP slopes inter-individually mirror model-based (HDDM) estimates of evidence integration (Figure S1-5c). Second, similar ramping dynamics are observed in the motor system (Text S1-6, central ERP above), where probe impact is more limited. Notably, contralateral beta power reflects relative biases towards one hemispheric response over the other, thereby fulfilling strict definitions of a choice-specific signature (Frömer et al., 2024). Effector-specific signatures (Donner et al., 2009) therefore lend further credence to the notion that our neural signatures capture evolving decision processes.

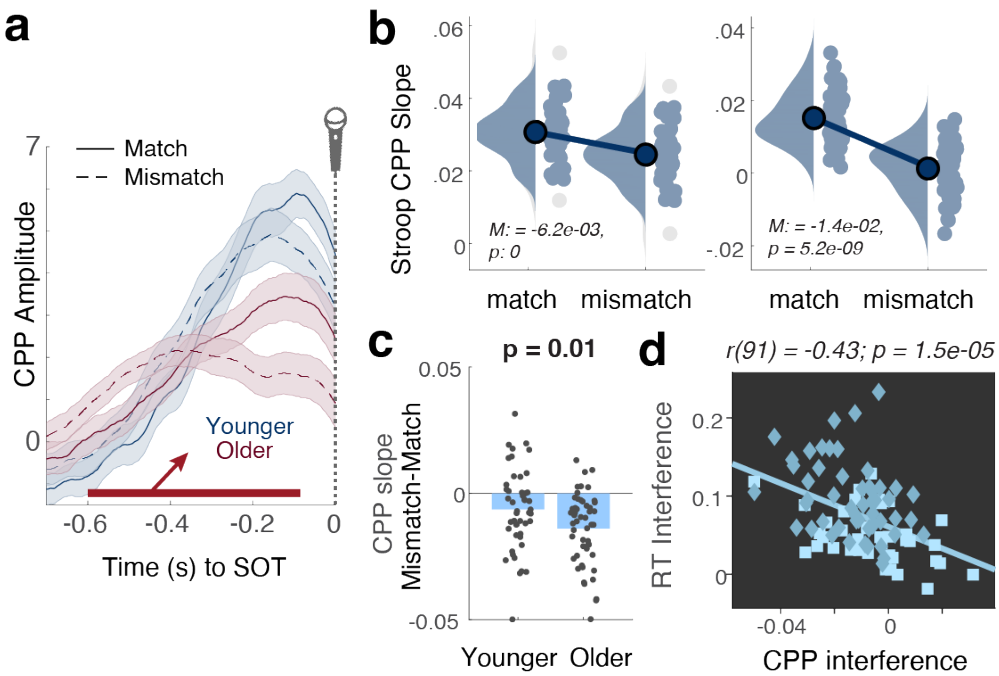


**Figure S3-1. CPP slope during the Stroop task.** (a) Response-aligned CPP traces split by condition and age group. Time series were smoothed with 60 ms windows for visualization, but not for slope fitting. Linear slopes were estimated during the interval of -600 to -100 ms prior to indicated SOTs, marked by the red line. (a) CPP integration slopes were reduced in magnitude in the mismatch condition in both age groups. (b) Interference effects on CPP slopes were more pronounced in younger adults compared with older adults. The magnitude of individual interference effects was similarly reflected in RTs and CPP slopes with longer RTs being coupled to stronger CPP slope reductions [r(91) = -0.43, 95%CI = [-0.58,-0.25], p = 1.5e-05; partial-correlation accounting for age: r(89) = -0.32, 95%CI = [-0.49,-0.12] p = 3.3e-04].

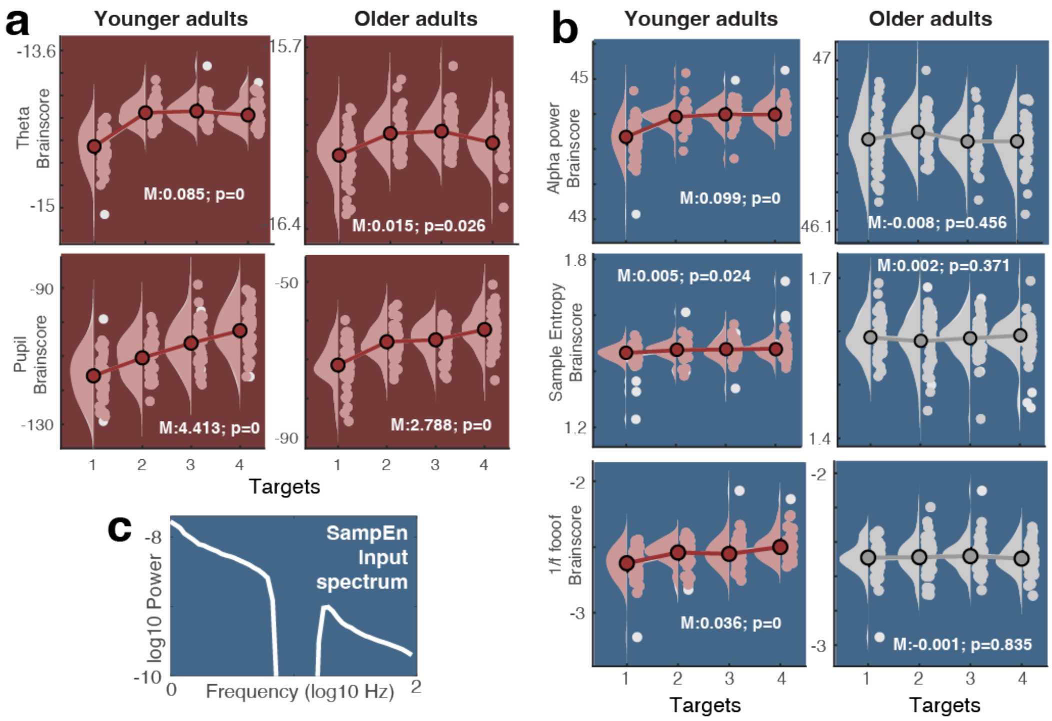


**Figure S4-1. Load modulation for cognitive control (a) and excitability signatures (b).** Statistics refer to paired t-tests of linear slopes against zero. In line with the different excitability indices capturing a shared latent characteristic, the magnitude of uncertainty modulation was inter-individually related among the three parameters (alpha-1/f: $r = 0.44$, $p = 9.8e-06$; 1/f-SampEn: $r = 0.6$, $p = 1.1e-10$; SampEn-alpha: $r = 0.24$, $p = .02$). (c) Sample entropy input spectrum highlighting the exclusion of alpha-range signal content.

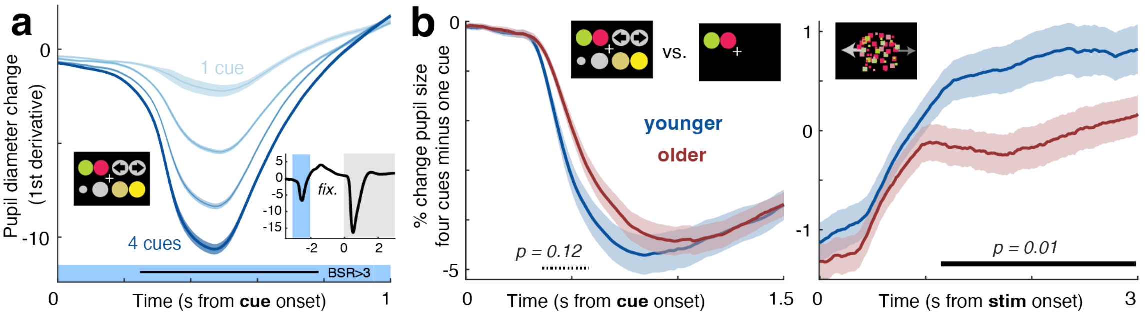


**Figure S4-2. Pupil size modulation in response to cue onset.** (a) Temporal derivatives of pupil diameter linearly decreased with increasing number of feature cues. Data are means +- within-subject SEM across age groups. The black line indicates significant loadings of a Task PLS LV1 (*permuted* p ~ 0). The inset shows grand average pupil dynamics with shaded cue and stimulus phases. (b) Relative pupil constriction upon presentation of more cues (= increasing luminance) did not differ between age groups, whereas younger adults showed larger relative pupil size for (luminance-matched) stimuli in the face of more uncertain feature relevance than older adults.

**Text S4-2. Stimulus-related pupil modulation does not result from cue-related luminance differences.** The task cue is composed of one to four bright exemplars. It thereby linearly increases in luminance alongside our uncertainty manipulation. To test whether luminance changes in the cueing phase affected stimulus-related pupil modulation, we probed pupil dynamics that were time-locked to cue onset (Figure S4-2a). Upon cue onset pupil size linearly decreased, in line with increasing luminance. These changes were however constrained to the cue presentation interval. Uncertainty-related pupil size changes upon stimulus presentation thus cannot be attributed to luminance differences in the preceding cue. Complementing the perspective on the 1$^{st}$ derivative of pupil dynamics, we also probed relative pupil size changes (calculated as % change relative to a condition-specific 500 ms baseline prior to cue onset: *data-baseline)/baseline*). Changes between max. and min. target load (= maximal cue luminance differences) are shown by age group in Fig. S4-2. Age groups did not significantly differ in relative pupil constriction upon cue presentation (~ luminance-effect; indicated by a cluster-based permutation test), but younger adults showed relatively increased pupil size in the face of more uncertain feature relevance following the onset of (luminance-matched) stimuli. These results jointly indicate that stimulus-related age differences in uncertainty-related pupil size modulation (shown in Fig. 4b) are not merely a spill-over of differential luminance sensitivity.

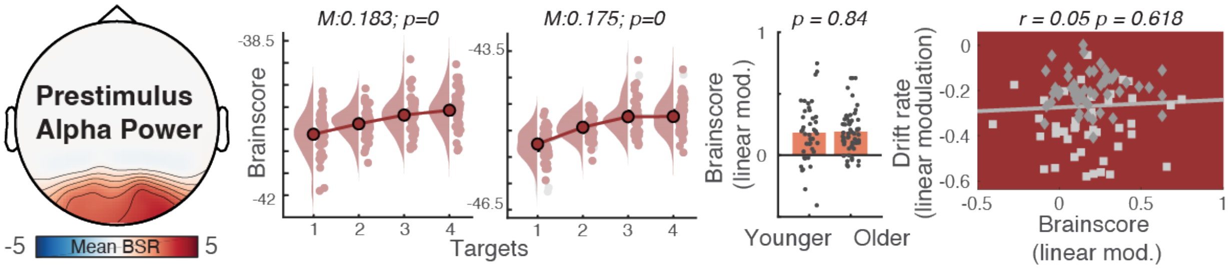


**Figure S5-1. Pre-stimulus alpha power.** Uncertainty similarly increases pre-stimulus alpha power in younger and older adults but does not relate to individual drift rate adjustments. Light grey squares indicate younger adults, dark grey diamonds correspond to older adults.

**Text S5-1 Pre-stimulus alpha power.** Evidence on age-related changes in pre-stimulus alpha power are mixed. Early studies suggest that pre-stimulus alpha synchronization (or lateralization) in the context of attentional cueing is observed exclusively for younger, but not older adults (Hong et al., 2015; Vaden et al., 2012). In contrast, (Leenders et al., 2018) indicated similar pre-stimulus lateralization between age groups, whereas they noted age differences in alpha modulation during working memory retention. While our task design does not allow us to assess the lateralization of alpha power, our results indicate that pre-stimulus alpha power increases similarly alongside uncertainty in both age groups, but with no apparent relation to subsequent (delayed) task performance (Figure S3-1).

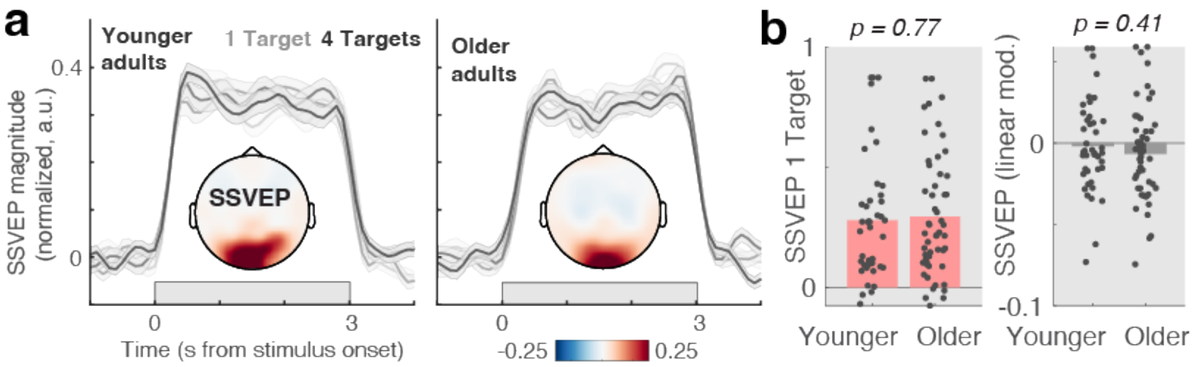


**Figure S5-2. Steady-state visual evoked potential (SSVEP).** (a) Both age groups exhibited a robust SSVEPs. Time-resolved, spectrally normalized, SSVEP power, averaged across occipital channels (O1, Oz, O2), indicates clear SSVEP increases specifically during stimulus presentation. Data are presented as mean values +/- within-subject SEM. Topography insets show stimulus-evoked SSVEP contrast minus baseline. (b) However, estimates from occipital EEG channels (O1, Oz, O2) did not indicate age differences in single-target SSVEP magnitude, a main effect of load in either group, or differences in the strength of linear modulation (~ age*load interaction).

**Text S5-2. SSVEP magnitude.** SSVEP magnitude has been suggested as a signature of encoded sensory information that is enhanced by attention (Morgan et al., 1996; Muller et al., 2006; Quigley et al., 2010; Quigley & Muller, 2014) and indicates fluctuations in early visual cortex excitability (Zhigalov et al., 2019). However, despite a clear SSVEP signature of comparable magnitude in both younger and older adults (Fig. S5-2 a, b), we did not observe significant effects of target uncertainty on SSVEP magnitude in either age group (Fig. S5-2). Given that the SSVEP frequency was shared across different features, we could not investigate feature selection via SSVEPs as is commonly the case in attention studies. Studies with feature-specific SSVEPs, suggest that younger adults' SSVEP magnitude differentiates between attended and unattended features, whereas no robust differentiation is observed in older adults, pointing to deficits in attentional filtering (Quigley et al., 2010; Quigley & Muller, 2014).

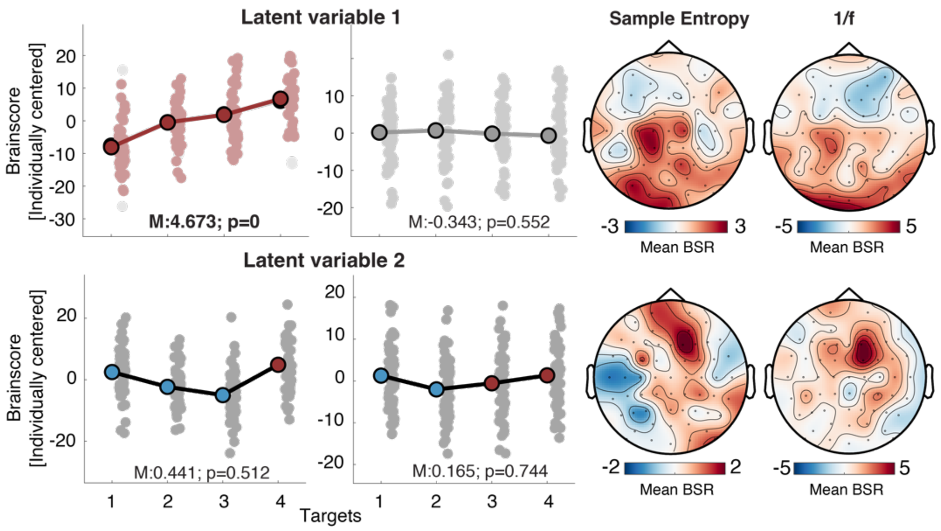


**Figure S5-3. Task PLS of sample entropy and aperiodic slopes across all channels.** Brainscores for younger adults are shown on the left side, with data for older adults shown to the right. Inset estimates refer to fixed linear effects models. Topographies of bootstrap ratios are unthresholded.

**Text S5-3. Exploratory whole-brain task PLS of aperiodic dynamics.** In the main analysis, we restricted the PLS to posterior channels with the aim to predominantly characterize signals stemming from parietal and visual cortex. To explore whether this analysis missed uncertainty-related changes in aperiodic dynamics in other regions, we performed an additional task PLS analysis that included all channels. This task PLS averaged sample entropy across the final 2.5s of stimulus presentation. To normalize relative contributions of the two signatures to the PLS, we z-transformed values of each signature across target load levels prior to including them in the model. This joint PLS resulted in two significant latent variables (Figure S5-3). The first latent variable (*permuted* p = 0.001) indicated uncertainty-related increases in sample entropy and shallowing of aperiodic slopes in younger, but not older adults. Regional contributions were predominantly observed in posterior sensors. This latent variable thus captures the observations in the main analysis. The second latent variable (*permuted* p = 0.021) was instead marked by quadratic changes (younger adults: p = 1.5e-08; older adults: p = 0.03; linear mixed effects model with fixed and random quadratic effects) as a function of target load. Estimates initially decreased, followed by an increase with load towards higher target load, predominantly at mediofrontal channels.

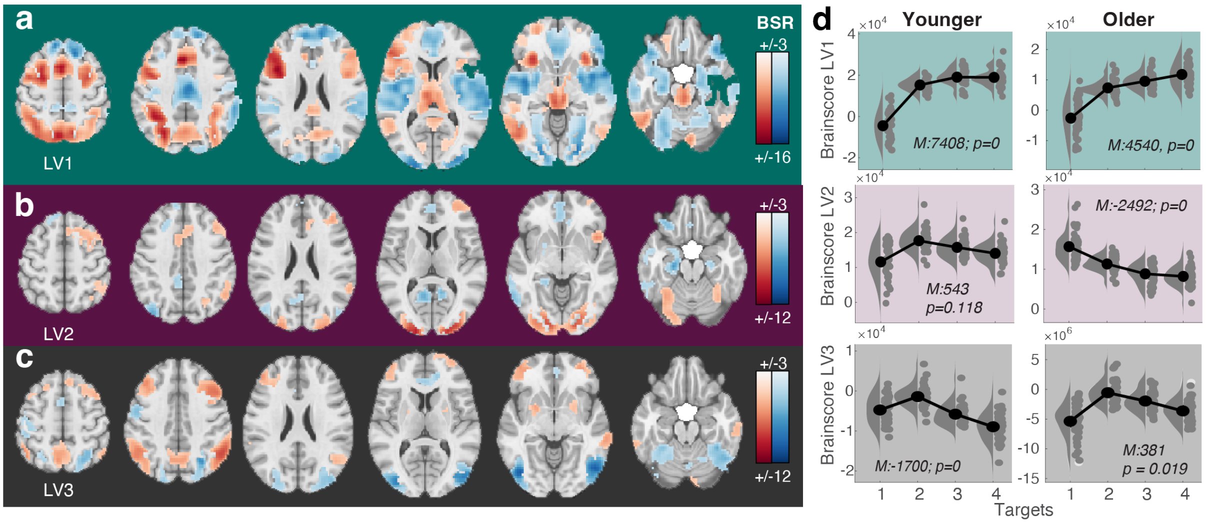


**Figure S6-1. Main effects of target load on BOLD magnitude.** A task partial least squares (PLS) analysis indicated three significant latent variables (loadings shown in panels a-c) that were sensitive to changes in target number. (d) Statistics refer to paired t-tests of linear slopes against zero.

**Text S6-1. Main effects of target load on BOLD magnitude across the adult lifespan.** We performed a whole-brain task PLS to assess potential main effects of target load on BOLD magnitude. In brief, we observed a similar first latent variable (*permuted* $p < 0.001$) to that reported in younger adults (Kosciessa et al., 2021), highlighting uncertainty-related increases dominantly in cortical areas encompassing the frontoparietal and the midcingulo-insular network, as well as in the thalamus (see detailed results of this analysis in Figure S6-1 and Table S2-4). The task PLS indicated two further robust LVs. LV2 (*permuted* $p < 0.001$) captured a non-linear pattern in younger adults and linear changes under uncertainty in older adults. Regional contributors partly overlapped with the initial LV (Table S3). Finally, LV3 (*permuted* $p < 0.001$) captured nonlinear changes (initial increases in engagement followed by disengagement) in both age groups in a set of regions encompassing positive loadings in frontoparietal components of the executive control network, and negative loadings in temporal-occipital cortex (Table S4).

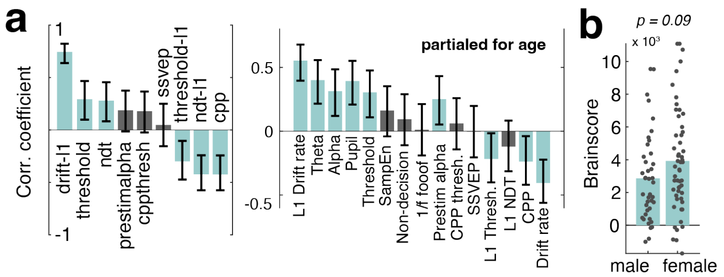


**Figure S6-2. Post-hoc Brainscore relations.** (a) Relations before (left) and after controlling for categorical age (right). Beyond the *a priori* signatures included in the behavioral PLS model, post-hoc exploration indicated that individuals with more pronounced BOLD uncertainty modulation also had larger single-target drift rates, and lower single-target boundary separation ("boundary thresholds"), as well as larger increases in the latter as a function of uncertainty, also after controlling for categorical age (see right). In addition to the magnitude of uncertainty-related drift rate modulation, CPP slope modulation was similarly related to Brainscores. Plots indicate Pearson correlation coefficients +- 95%CI after accounting for age covariation. (b) Brainscores from behavioral PLS as a function of gender.

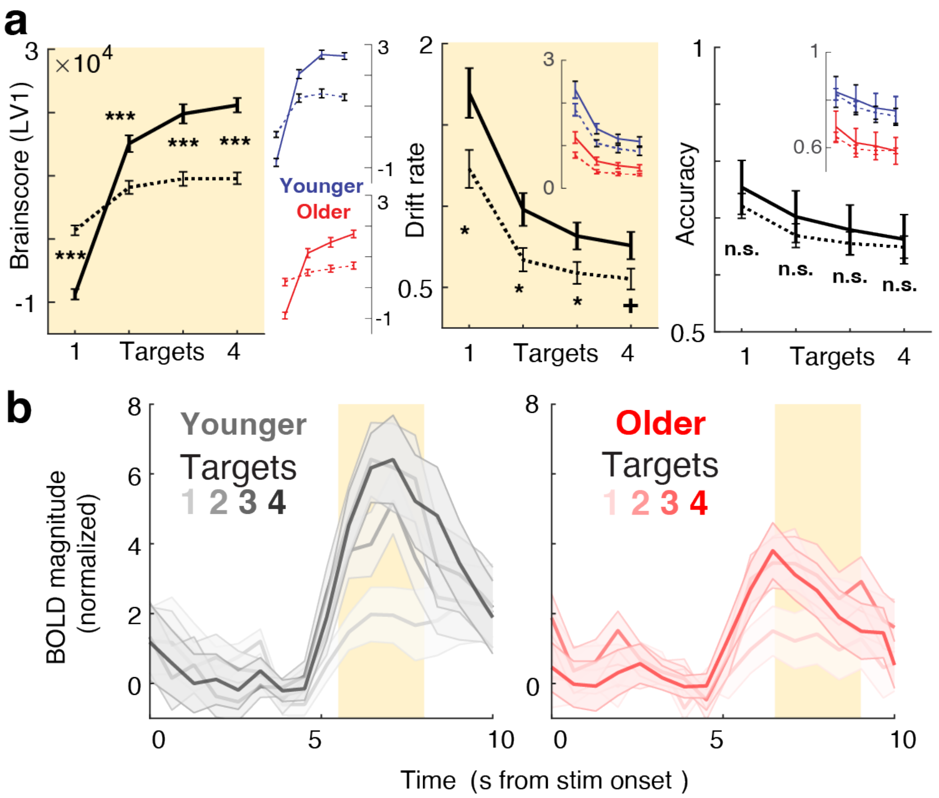


**Figure S6-3. BOLD modulation effects are robust to accuracy differences within (a) and between age groups (b).** (a) Younger and older individuals with larger BOLD uncertainty modulation (LV1, see Fig S6-1) achieve higher drift rates across uncertainty levels at comparable accuracy levels. Data show upper (full lines) and lower (broken lines) groups of a trichotomized split of accuracy and drift rate data based on the magnitude of uncertainty change (234 vs. 1) in the 1$^{st}$ LV of the task PLS (closely mirroring LV1 of the behavioral PLS). Insets illustrate comparable within-group splits (split performed within-group). Data are means +- SEs. (b) Age x uncertainty interaction in mediodorsal thalamus for accuracy-matched features. This analysis excluded trials in which the best (YA) or worst (OA) features were probed to compare data with group-matched single-target accuracy (see Text S1-3). A linear mixed effects model indicated a retained group x target load interaction for data averaged in the time window of interest (yellow shading; beta=-0.757, SE=0.369, t = -2.0507, dof = 364, p = 0.041, 95%CI = [-1.48, -0.03]). Data are means +- SEs.

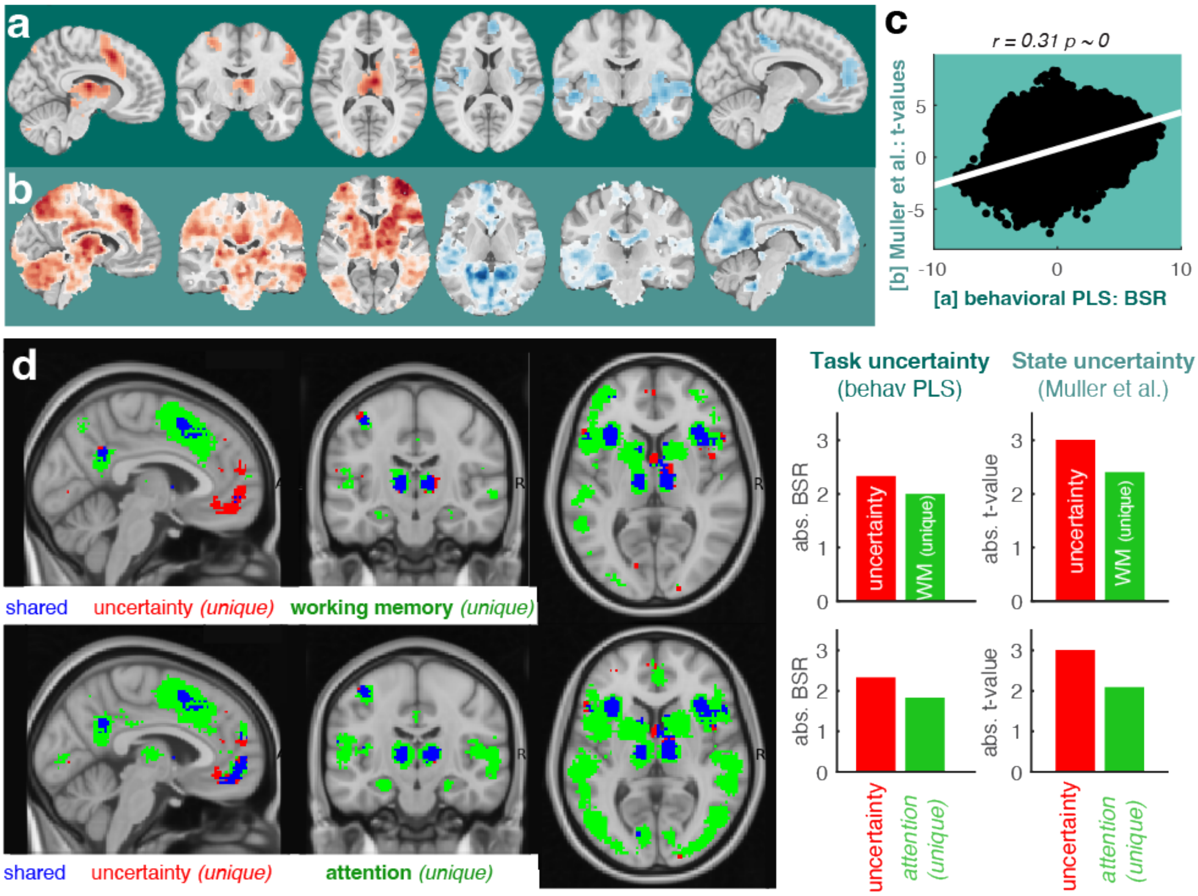


**Figure S6-4. Reverse inference of uncertainty, working memory, and attention.** (a-c) Overlap between neuroimaging of task and state uncertainty. (a) Spatial loadings of behavioral PLS, reproduced from Fig. 6a. (b) Neuroimaging results of a state entropy manipulation with low working memory demands [reanalyzed from Fig. 1-3a in (Muller et al., 2019); https://neurovault.org/collections/4872/]. Data range is [0 to -/+ 6]. (c) Person correlation between spatial t-values and spatial loadings indicate convergent spatial activation patterns between uncertainty manipulations. (d) Overlap and uniqueness of uncertainty and working memory activation. Neurosynth.org activations (uniformity tests: the degree to which each voxel is consistently activated in studies that use a given term; threshold: FDR = 0.01; binarization threshold: 1) for "working memory" and "uncertainty", decomposed into shared and unique patterns. Both the current task uncertainty modulation and state uncertainty more closely resemble meta-analytic loadings of uncertainty than working memory or attention patterns.

**Text S6-4. Relation of fMRI modulation to uncertainty, working memory, and attention.** Our uncertainty manipulation demands multifaceted adjustments, including to attentional selection, extra-dimensional set switching and working memory maintenance. Can our results be directly linked to a single cognitive concept? We performed reverse inference analyses to clarify this. In line with our results capturing variations in induced uncertainty, neuroimaging results converged with a prior assessment of model-based uncertainty (about the reward probability of eight options in an Bayesian ideal observer model (Muller et al., 2019); **Fig. S6-4a-c**). In this prior operationalization, uncertainty varies according to varying probabilistic beliefs across features, but does not directly scale with working memory load. Our results converge with raw loadings from this prior assessment (incl. on thalamic loadings not originally highlighted). When directly contrasting task uncertainty activation to meta-analytic activations to attention or working memory studies (see **Fig. S6-4d**), neuroimaging results were more consistent with meta-analytic maps of uncertainty. This was also true when considering % overlap between binary empirical activation (+/-1.96 threshold) and neurosynth masks: task uncertainty: uncertainty: 53% | wm: 47% | attention: 41%; state uncertainty: uncertainty: 78% | wm: 64%| attention: 53%. Taken together, these results indicate that our results converge with prior work on uncertainty, while also indicating overlap with multifaceted executive adjustment in response to variable uncertainty.

**Text S7. Extended discussion: uncertainty and working memory capacity.** Can decreasing working memory (WM) capacity with older age (Naveh-Benjamin & Cowan, 2023) account for dampened uncertainty adjustment? Our task consists of three stages, each of which requires WM representations. First, from cue to probe presentation, participants need to maintain 1 to 4 targets in the active task set. We aimed to reduce WM load during this phase by maintaining an identical cue set within eight-trial blocks, and re-cueing this set at the onset of each trial. Cue-related EEG responses indicate similar alpha power modulation for younger and older adults (Fig. S5-1), suggesting comparable processing of the cued feature set and alertness. Second, during stimulus presentation, the MAAT requires participants to concurrently sample up to four visual features and retain WM traces of feature-specific information for the upcoming probe-dependent decision phase. Third, once a probe retro-cues the decision-relevant feature, feature-specific evidence must be selected amongst competing WM representations (Mok et al., 2016). Probe processing (and behaviour) can be affected by how many features could be encoded given individual WM capacity across latter task stages. However, behavioural (and neural) uncertainty effects (and age differences therein) were prominent also at low target loads, arguing against a major role of WM capacity constraints. Rather, the pattern of effects supports age differences in executive control over WM contents, as also observed in N-back tasks (Bopp & Verhaeghen, 2020). A capacity-based explanation is also less intuitive for uncertainty-induced changes at the onset of stimulus presentation (e.g., increased pupil diameter, EEG-based power). We argue that WM *fidelity* is more important in this context than WM *capacity* per se (Hill et al., 2021). In line with this notion, our decoding analysis indicates an increase in information about multiple features at the expense of single-target fidelity. Operationalizations of task uncertainty via cue conflict (as operationalized here; see also (Mukherjee et al., 2021; Tsumura et al., 2022)) are also used to probe WM fidelity under varying target predictability. Studies of the latter indicate that uncertainty reduces the precision of (probabilistic) WM representations (Li et al., 2021; Ma et al., 2014), here theoretically for each feature (the feature-specific states shown in Fig. 7). Neuroimaging further supports a close overlap between uncertainty and WM load manipulations (see also Fig. S6-4): sensitivity to either relies on a "multiple demand network" (Duncan, 2010). Notably, mediodorsal thalamic engagement has been theorized to be maximal when multiple demands (incl. executive control operations such as WM updating, inhibition, and shifting) converge (Pergola et al., 2018). Our results therefore tacitly support the view that older adults are particularly challenged in dynamic WM management (Zuber et al., 2019), and related parallel resource allocation (Jaroslawska et al., 2021), whereas WM storage capacity shows only modest age-related declines (for review see Naveh-Benjamin & Cowan, 2023).

**Table S1: Statistics for within age-group effects.** Effects were assessed via paired t-tests against zero.

| Dependent variable | Fig. | df | t-value | p-value | Cohen's d |
|---|---|---|---|---|---|
| Drift rate (single-target) – younger | 1b | 41 | 26.41 | 2.3e-27 | 4.07 |
| Drift rate (single-target) – older | 1b | 52 | 22.06 | 4.6e-28 | 3.03 |
| CPP (single-target) – younger | 1b | 41 | 8.62 | 9.5e-11 | 1.33 |
| CPP (single-target) – older | 1b | 52 | 7.92 | 1.7e-10 | 1.09 |
| Drift rate (linear mod.) – younger | 1b | 41 | -17.07 | 3.1e-20 | -2.63 |
| Drift rate (linear mod.) – older | 1b | 52 | -17.45 | 2.2e-23 | -2.4 |
| CPP (linear mod.) – younger | 1b | 41 | -7.37 | 4.9e-09 | -1.14 |
| CPP (linear mod.) – older | 1b | 52 | -5.04 | 5.9e-06 | -0.69 |
| Stroop interference – younger | 3b | 47 | 10.01 | 3.1e-13 | 1.44 |
| Stroop interference – older | 3b | 52 | 16.02 | 9.3e-22 | 2.2 |
| Theta power (linear mod.) – younger | 4a | 41 | 6.85 | 2.6e-08 | 1.06 |
| Theta power (linear mod.) – older | 4a | 52 | 2.3 | 2.6e-02 | 0.32 |
| Pupil diameter (linear mod.) – younger | 4b | 41 | 7.34 | 5.6e-09 | 1.13 |
| Pupil diameter (linear mod.) – older | 4b | 52 | 7.25 | 1.9e-09 | 1 |
| Alpha power (linear mod.) – younger | 5a | 41 | 6.05 | 3.7e-07 | 0.93 |
| Alpha power (linear mod.) – older | 5a | 52 | -0.75 | 0.46 | -0.1 |
| Sample Entropy (linear mod.) – younger | 5b | 41 | 2.21 | 0.033 | 0.34 |
| Sample Entropy (linear mod.) – older | 5b | 52 | 0.23 | 0.82 | 0.03 |
| 1/f slope (linear mod.) – younger | 5c | 41 | 4.67 | 3.3e-05 | 0.72 |
| 1/f slope (linear mod.) – older | 5c | 52 | 0.14 | 0.89 | 0.02 |
| fMRI Brainscore – younger | 6b | 41 | 11.49 | 2.1e-14 | 1.77 |
| fMRI Brainscore – older | 6b | 52 | 7.47 | 8.8e-10 | 1.03 |

**Table S2: PLS model peak activations, bootstrap ratios, and cluster sizes for task PLS LV1.**

| | | MNI Coordinates | | | | |
|---|---|---|---|---|---|---|
| Region | Hem | X | Y | Z | BSR | #Voxels |
| IFG (p. Opercularis) | L | -45 | 9 | 27 | 15.11 | 3164 |
| Inferior Parietal Lobule | L | -42 | -48 | 45 | 14.3 | 3451 |
| Insula Lobe | R | 30 | 21 | -3 | 11.4 | 170 |
| Inferior Temporal Gyrus | L | -54 | -66 | -12 | 11.34 | 880 |
| Thalamus | L | -6 | -30 | -3 | 10.76 | 1064 |
| Superior Frontal Gyrus | R | 27 | -3 | 54 | 10.21 | 903 |
| Cerebelum (Crus 1) | R | 6 | -81 | -24 | 8.5 | 276 |
| Cerebelum (VI) | R | 30 | -66 | -30 | 7.83 | 129 |
| Inferior Temporal Gyrus | R | 54 | -63 | -12 | 6.19 | 297 |
| Area Fo3 | L | -27 | 39 | -21 | 4.68 | 45 |
| Calcarine Gyrus | L | -15 | -78 | 6 | 4.28 | 29 |
| Middle Frontal Gyrus | R | 27 | 51 | 3 | 4.25 | 31 |
| Superior Medial Gyrus | R | 12 | 48 | 33 | -12.32 | 2317 |
| Area hOc3d [V3d] | L | -24 | -99 | 12 | -11.64 | 6542 |
| MCC | R | 3 | -15 | 36 | -11.23 | 889 |
| Area lg1 | R | 30 | -21 | 3 | -11.14 | 4238 |
| Postcentral Gyrus | R | 21 | -36 | 63 | -5.86 | 121 |
| Postcentral Gyrus | L | -39 | -21 | 36 | -5.8 | 43 |
| Middle Frontal Gyrus | L | -33 | 24 | 39 | -5.32 | 32 |
| Angular Gyrus | L | -48 | -63 | 27 | -4.81 | 59 |
| Middle Frontal Gyrus | R | 45 | 15 | 48 | -3.95 | 55 |

**Table S3: PLS model peak activations, bootstrap ratios, and cluster sizes for task PLS LV2.**

| | | MNI Coordinates | | | | |
|---|---|---|---|---|---|---|
| Region | Hem | X | Y | Z | BSR | #Voxels |
| Area hOc3d [V3d] | L | -24 | -99 | 12 | 9.16 | 1974 |
| Insula Lobe | R | 42 | 15 | -3 | 6.66 | 136 |
| Middle Orbital Gyrus | R | 30 | 54 | -15 | 5.31 | 143 |
| Superior Frontal Gyrus | R | 21 | 12 | 60 | 5.01 | 911 |
| Middle Frontal Gyrus | R | 33 | 48 | 12 | 4.91 | 119 |
| Angular Gyrus | R | 57 | -51 | 27 | 4.76 | 280 |
| Inferior Temporal Gyrus | R | 51 | -3 | -39 | 4.47 | 50 |
| Inferior Temporal Gyrus | R | 63 | -27 | -30 | 4.4 | 28 |
| Superior Occipital Gyrus | R | 21 | -63 | 42 | 4.05 | 69 |
| Rolandic Operculum | L | -57 | 6 | 3 | 3.79 | 27 |
| Hippocampus | L | -27 | -21 | -18 | -7.36 | 316 |
| Calcarine Gyrus | L | -12 | -60 | 12 | -6.5 | 244 |
| Rectal Gyrus | L | -9 | 27 | -15 | -6.24 | 440 |
| Middle Temporal Gyrus | L | -66 | -57 | -9 | -6.12 | 256 |
| Middle Occipital Gyrus | L | -42 | -81 | 39 | -6.08 | 190 |
| IFG (p. Orbitalis) | L | -36 | 33 | -18 | -5.97 | 94 |
| Precuneus | R | 9 | -54 | 9 | -5.67 | 131 |
| ParaHippocampal Gyrus | R | 21 | -21 | -18 | -4.83 | 39 |
| IFG (p. Orbitalis) | R | 24 | 27 | -15 | -4.67 | 28 |
| Middle Temporal Gyrus | R | 54 | -6 | -15 | -4.56 | 51 |
| MCC | L | -12 | -42 | 36 | -4.44 | 76 |
| Middle Occipital Gyrus | R | 45 | -78 | 27 | -4.34 | 34 |
| Middle Frontal Gyrus | L | -27 | 30 | 42 | -4.27 | 153 |
| Cerebelum (Crus 2) | R | 3 | -87 | -33 | -4.14 | 28 |
| Middle Temporal Gyrus | L | -51 | -3 | -24 | -3.94 | 53 |
| Superior Frontal Gyrus | L | -24 | 60 | 3 | -3.78 | 27 |

**Table S4: PLS model peak activations, bootstrap ratios, and cluster sizes for task PLS LV3.**

| | | MNI Coordinates | | | | |
|---|---|---|---|---|---|---|
| Region | Hem | X | Y | Z | BSR | #Voxels |
| Angular Gyrus | R | 51 | -57 | 36 | 8.05 | 604 |
| Middle Frontal Gyrus | R | 36 | 21 | 36 | 6.87 | 660 |
| Inferior Parietal Lobule | L | -54 | -51 | 42 | 6.27 | 469 |
| Precuneus | L | -9 | -66 | 45 | 6.13 | 474 |
| Middle Frontal Gyrus | L | -39 | 24 | 33 | 6.05 | 726 |
| Middle Frontal Gyrus | R | 27 | 57 | 0 | 5.97 | 287 |
| Middle Temporal Gyrus | R | 60 | -33 | -12 | 5.46 | 184 |
| Cerebelum (Crus 1) | R | 9 | -81 | -27 | 5.29 | 62 |
| Putamen | L | -27 | 6 | -6 | 4.82 | 74 |
| Putamen | R | 24 | 0 | 6 | 4.38 | 67 |
| Inferior Temporal Gyrus | L | -66 | -42 | -21 | 4.16 | 49 |
| Cerebelum (Crus 2) | L | -15 | -87 | -30 | 3.96 | 32 |
| Cerebelum (Crus 2) | R | 33 | -72 | -45 | 3.8 | 53 |
| Inferior Temporal Gyrus | R | 48 | -69 | -9 | -10.33 | 1706 |
| Inferior Occipital Gyrus | L | -45 | -75 | -6 | -9.9 | 1022 |
| Postcentral Gyrus | L | -57 | -6 | 39 | -5.18 | 232 |
| Postcentral Gyrus | L | -51 | -33 | 57 | -4.5 | 43 |
| ACC | R | 12 | 42 | 9 | -4.41 | 191 |
| Superior Parietal Lobule | L | -24 | -63 | 48 | -4.4 | 36 |
| Posterior-Medial Frontal | L | -6 | 3 | 57 | -4.4 | 65 |

**Table S5: PLS model peak activations, bootstrap ratios, and cluster sizes for behavioral PLS LV1.**

| | | MNI Coordinates | | | | |
|---|---|---|---|---|---|---|
| Region | Hem | X | Y | Z | BSR | #Voxels |
| Thalamus | L | -6 | -15 | 12 | 8.57 | 573 |
| Posterior-Medial Frontal | L | 3 | 12 | 45 | 8.13 | 555 |
| Precentral Gyrus | L | -42 | 0 | 30 | 7.51 | 931 |
| Superior Frontal Gyrus | R | 27 | -3 | 54 | 6.91 | 222 |
| Inferior Occipital Gyrus | L | -42 | -72 | -6 | 6.13 | 208 |
| Middle Temporal Gyrus | L | -51 | -51 | 21 | 6.03 | 90 |
| Putamen | R | 30 | 18 | 0 | 5.75 | 94 |
| Middle Frontal Gyrus | R | 36 | 24 | 21 | 5.74 | 220 |
| Middle Temporal Gyrus | L | -57 | -33 | -6 | 5.26 | 35 |
| Inferior Parietal Lobule | R | 30 | -54 | 48 | 5.14 | 323 |
| Inferior Parietal Lobule | L | -36 | -57 | 45 | 4.75 | 315 |
| Area hOc1 [V1] | R | 9 | -99 | 6 | 4.7 | 82 |
| Inferior Temporal Gyrus | R | 45 | -63 | -12 | 4.51 | 27 |
| Hippocampus | L | -27 | -18 | -21 | -8.92 | 873 |
| MCC | L | -12 | -36 | 48 | -6.05 | 359 |
| Putamen | R | 30 | -3 | 9 | -5.79 | 879 |
| Superior Frontal Gyrus | R | 18 | 54 | 30 | -5.73 | 443 |
| Middle Frontal Gyrus | L | -21 | 30 | 54 | -5.45 | 67 |
| Superior Medial Gyrus | R | 12 | 42 | 48 | -5.28 | 138 |
| IFG (p. Orbitalis) | R | 36 | 30 | -21 | -5.2 | 55 |
| ParaHippocampal Gyrus | R | 21 | -18 | -18 | -5.13 | 49 |
| Middle Temporal Gyrus | L | -51 | 3 | -33 | -4.28 | 29 |
| Inferior Frontal Gyrus | L | -33 | 36 | -21 | -4.17 | 26 |
| Rectal Gyrus | L | -9 | 24 | -12 | -4.15 | 73 |
| Inferior Temporal Gyrus | L | -57 | -24 | -27 | -4.03 | 32 |

### Supplementary References


Bopp, K. L., & Verhaeghen, P. (2020). Aging and n-Back Performance: A Meta-Analysis. *J Gerontol B Psychol Sci Soc Sci*, *75*(2), 229-240. https://doi.org/10.1093/geronb/gby024

Churchland, A. K., Kiani, R., & Shadlen, M. N. (2008). Decision-making with multiple alternatives. *Nat Neurosci*, *11*(6), 693-702. https://doi.org/10.1038/nn.2123

Donner, T. H., Siegel, M., Fries, P., & Engel, A. K. (2009). Buildup of choice-predictive activity in human motor cortex during perceptual decision making. *Curr Biol*, *19*(18), 1581-1585. https://doi.org/10.1016/j.cub.2009.07.066

Duncan, J. (2010). The multiple-demand (MD) system of the primate brain: mental programs for intelligent behaviour. *Trends Cogn Sci*, *14*(4), 172-179. https://doi.org/10.1016/j.tics.2010.01.004

Ehinger, B. V., & Dimigen, O. (2019). Unfold: an integrated toolbox for overlap correction, non-linear modeling, and regression-based EEG analysis. *PeerJ*, *7*, e7838. https://doi.org/10.7717/peerj.7838

Frömer, R., Nassar, M. R., Ehinger, B. V., & Shenhav, A. (2024). Common neural choice signals can emerge artifactually amidst multiple distinct value signals. *bioRxiv*, 2022.2008.2002.502393. https://doi.org/10.1101/2022.08.02.502393

Hill, P. F., King, D. R., & Rugg, M. D. (2021). Age Differences In Retrieval-Related Reinstatement Reflect Age-Related Dedifferentiation At Encoding. *Cereb Cortex*, *31*(1), 106-122. https://doi.org/10.1093/cercor/bhaa210

Hong, X. F., Sun, J. F., Bengson, J. J., Mangun, G. R., & Tong, S. B. (2015). Normal aging selectively diminishes alpha lateralization in visual spatial attention. *NeuroImage*, *106*, 353-363. https://doi.org/10.1016/j.neuroimage.2014.11.019

Jaroslawska, A. J., Rhodes, S., Belletier, C., Doherty, J. M., Cowan, N., Neveh-Benjamin, M., Barrouillet, P., Camos, V., & Logie, R. H. (2021). What affects the magnitude of age-related dual-task costs in working memory? The role of stimulus domain and access to semantic representations. *Q J Exp Psychol (Hove)*, *74*(4), 682-704. https://doi.org/10.1177/1747021820970744

Kosciessa, J. Q., Lindenberger, U., & Garrett, D. D. (2021). Thalamocortical excitability modulation guides human perception under uncertainty. *Nature Communications*, *12*(1), 2430. https://doi.org/10.1038/s41467-021-22511-7

Leenders, M. P., Lozano-Soldevilla, D., Roberts, M. J., Jensen, O., & De Weerd, P. (2018). Diminished Alpha Lateralization During Working Memory but Not During Attentional Cueing in Older Adults. *Cereb Cortex*, *28*(1), 21-32. https://doi.org/10.1093/cercor/bhw345

Li, H. H., Sprague, T. C., Yoo, A. H., Ma, W. J., & Curtis, C. E. (2021). Joint representation of working memory and uncertainty in human cortex. *Neuron*, *109*(22), 3699-3712 e3696. https://doi.org/10.1016/j.neuron.2021.08.022

Ma, W. J., Husain, M., & Bays, P. M. (2014). Changing concepts of working memory. *Nat Neurosci*, *17*(3), 347-356. https://doi.org/10.1038/nn.3655

McGovern, D. P., Hayes, A., Kelly, S. P., & O'Connell, R. G. (2018). Reconciling age-related changes in behavioural and neural indices of human perceptual decision-making. *Nat Hum Behav*, *2*(12), 955-966. https://doi.org/10.1038/s41562-018-0465-6

Mok, R. M., Myers, N. E., Wallis, G., & Nobre, A. C. (2016). Behavioral and Neural Markers of Flexible Attention over Working Memory in Aging. *Cereb Cortex*, *26*(4), 1831-1842. https://doi.org/10.1093/cercor/bhw011

Morgan, S. T., Hansen, J. C., & Hillyard, S. A. (1996). Selective attention to stimulus location modulates the steady-state visual evoked potential. *Proc Natl Acad Sci U S A*, *93*(10), 4770-4774. https://doi.org/10.1073/pnas.93.10.4770

Mukherjee, A., Lam, N. H., Wimmer, R. D., & Halassa, M. M. (2021). Thalamic circuits for independent control of prefrontal signal and noise. *Nature*. https://doi.org/10.1038/s41586-021-04056-3

Muller, M. M., Andersen, S., Trujillo, N. J., Valdes-Sosa, P., Malinowski, P., & Hillyard, S. A. (2006). Feature-selective attention enhances color signals in early visual areas of the human brain. *Proc Natl Acad Sci U S A*, *103*(38), 14250-14254. https://doi.org/10.1073/pnas.0606668103

Muller, T. H., Mars, R. B., Behrens, T. E., & O'Reilly, J. X. (2019). Control of entropy in neural models of environmental state. *Elife*, *8*. https://doi.org/10.7554/eLife.39404

Naveh-Benjamin, M., & Cowan, N. (2023). The roles of attention, executive function and knowledge in cognitive ageing of working memory. *Nature Reviews Psychology*, *2*(3), 151-165. https://doi.org/10.1038/s44159-023-00149-0

O'Connell, R. G., Dockree, P. M., & Kelly, S. P. (2012). A supramodal accumulation-to-bound signal that determines perceptual decisions in humans. *Nat Neurosci*, *15*(12), 1729-1735. https://doi.org/10.1038/nn.3248

Pergola, G., Danet, L., Pitel, A. L., Carlesimo, G. A., Segobin, S., Pariente, J., Suchan, B., Mitchell, A. S., & Barbeau, E. J. (2018). The Regulatory Role of the Human Mediodorsal Thalamus. *Trends Cogn Sci*, *22*(11), 1011-1025. https://doi.org/10.1016/j.tics.2018.08.006

Quigley, C., Andersen, S. K., Schulze, L., Grunwald, M., & Muller, M. M. (2010). Feature-selective attention: evidence for a decline in old age. *Neurosci Lett*, *474*(1), 5-8. https://doi.org/10.1016/j.neulet.2010.02.053

Quigley, C., & Muller, M. M. (2014). Feature-selective attention in healthy old age: a selective decline in selective attention? *J Neurosci*, *34*(7), 2471-2476. https://doi.org/10.1523/JNEUROSCI.2718-13.2014

Starns, J. J., & Ratcliff, R. (2010). The effects of aging on the speed-accuracy compromise: Boundary optimality in the diffusion model. *Psychol Aging*, *25*(2), 377-390. https://doi.org/10.1037/a0018022
Starns, J. J., & Ratcliff, R. (2012). Age-related differences in diffusion model boundary optimality with both trial-limited and time-limited tasks. *Psychon Bull Rev*, *19*(1), 139-145. https://doi.org/10.3758/s13423-011-0189-3
Tsumura, K., Kosugi, K., Hattori, Y., Aoki, R., Takeda, M., Chikazoe, J., Nakahara, K., & Jimura, K. (2022). Reversible Fronto-occipitotemporal Signaling Complements Task Encoding and Switching under Ambiguous Cues. *Cereb Cortex*, *32*(9), 1911-1931. https://doi.org/10.1093/cercor/bhab324
Twomey, D. M., Murphy, P. R., Kelly, S. P., & O'Connell, R. G. (2015). The classic P300 encodes a build-to-threshold decision variable. *Eur J Neurosci*, *42*(1), 1636-1643. https://doi.org/10.1111/ejn.12936
Vaden, R. J., Hutcheson, N. L., McCollum, L. A., Kentros, J., & Visscher, K. M. (2012). Older adults, unlike younger adults, do not modulate alpha power to suppress irrelevant information. *NeuroImage*, *63*(3), 1127-1133. https://doi.org/10.1016/j.neuroimage.2012.07.050
Zhigalov, A., Herring, J. D., Herpers, J., Bergmann, T. O., & Jensen, O. (2019). Probing cortical excitability using rapid frequency tagging. *NeuroImage*, *195*, 59-66. https://doi.org/10.1016/j.neuroimage.2019.03.056
Zuber, S., Ihle, A., Loaiza, V. M., Schnitzspahn, K. M., Stahl, C., Phillips, L. H., Kaller, C. P., & Kliegel, M. (2019). Explaining age differences in working memory: The role of updating, inhibition, and shifting. *Psychology & Neuroscience*, *12*(2), 191-208. https://doi.org/10.1037/pne0000151
